# Dynamical networks: finding, measuring, and tracking neural population activity using network science

**DOI:** 10.1101/115485

**Authors:** Mark D. Humphries

## Abstract

Systems neuroscience is in a head-long rush to record from as many neurons at the same time as possible. As the brain computes and codes using neuron populations, it is hoped these data will uncover the fundamentals of neural computation. But with hundreds, thousands, or more simultaneously recorded neurons comes the inescapable problems of visualising, describing, and quantifying their interactions. Here I argue that network science provides a set of scalable, analytical tools that already solve these problems. By treating neurons as nodes and their interactions as links, a single network can visualise and describe an arbitrarily large recording. I show that with this description we can quantify the effects of manipulating a neural circuit, track changes in population dynamics over time, and quantitatively define theoretical concepts of neural populations such as cell assemblies. Using network science as a core part of analysing population recordings will thus provide both qualitative and quantitative advances to our understanding of neural computation.

---

Neurons use spikes to communicate (Rieke, Warland, de Ruyter van Stevninck, & Bialek, 1999). From this communication arises coding and computation within the brain; and so arises all thought, perception, and deed. Understanding neural circuits thus hinges critically on understanding spikes across populations of neurons (Pouget, Beck, Ma, & Latham, 2013; Wohrer, Humphries, & Machens, 2013; Yuste, 2015).

This idea has driven a technological arms race in systems neuroscience to record from as many individual neurons at the same time as physically possible (Stevenson & Kording, 2011). Current technology, ranging from imaging of fluorescent calcium-binding proteins (Chen et al., 2013; Dupre & Yuste, 2017; S. Peron, Chen, & Svoboda, 2015; S. P. Peron, Freeman, Iyer, Guo, & Svoboda, 2015) and voltage-sensitive dyes (Briggman, Abarbanel, & Kristan, 2005; Bruno, Frost, & Humphries, 2015; Frady, Kapoor, Horvitz, & Kristan, 2016) to large scale multi-electrode arrays and silicon probes (Buzsáki, 2004; Jun et al., 2017), now allows us to simultaneously capture the activity of hundreds of neurons in a range of brain systems. These systems include such diverse systems as invertebrate locomotion, through zebrafish oculomotor control, to executive functions in primate prefrontal cortex. With the data captured, the key question for any system becomes: how do we describe these spike data? Visualise them? And how do we discover the coding and computations therein?

Here I argue that network science provides a set of tools ideally suited to both describe the data and discover new ideas within it. Networks are simply a collection of nodes and links: nodes representing objects, and links representing the interactions between those objects. This representation can encapsulate a wide array of systems, from email traffic within a company, through the social groups of dolphins, to word co-occurrence frequencies in a novel (Newman, 2003). By abstracting these complex systems to a network description, we can describe their topology, compare them, and deconstruct them into their component parts. Moreover, we gain access to a range of null models for testing hypotheses about a network’s structure and about how it changes. I will demonstrate all these ideas below.

First, an important distinction. Networks capture interactions as links, but these links do not necessarily imply physical connections. In some cases, such as the network of router-level connections of the Internet or a power-grid, the interaction network follows exactly a physical network. In somes cases, such as a Facebook social network, there is no physical connection between the nodes. In other cases, of which neuroscience is a prime example, the interactions between nodes are shaped and constrained by the underlying physical connections, but are not bound to them. We shall touch on this issue of distinguishing interactions from physical connections throughout.

## DESCRIBING MULTI-NEURON DATA AS A NETWORK

A network description of multi-neuron recording data rests on two ideas: the nodes are the neurons, and the links are the interactions between the neurons (Figure 1A). Strictly speaking, the nodes are the isolated time-series of neural activity, whether spike-trains, calcium fluorescence, or voltage-dye expression (with the usual caveats applied to the accuracy of spike-sorting for electrodes or image segmentation and stability for imaging; Harris, Quiroga, Freeman, & Smith, 2016). An immediate advantage of a network formalism is that it separates the details of choosing the interaction from the network topology itself - whatever measure of interaction we chose, the same topological analyses can be applied.

**Figure 1.**
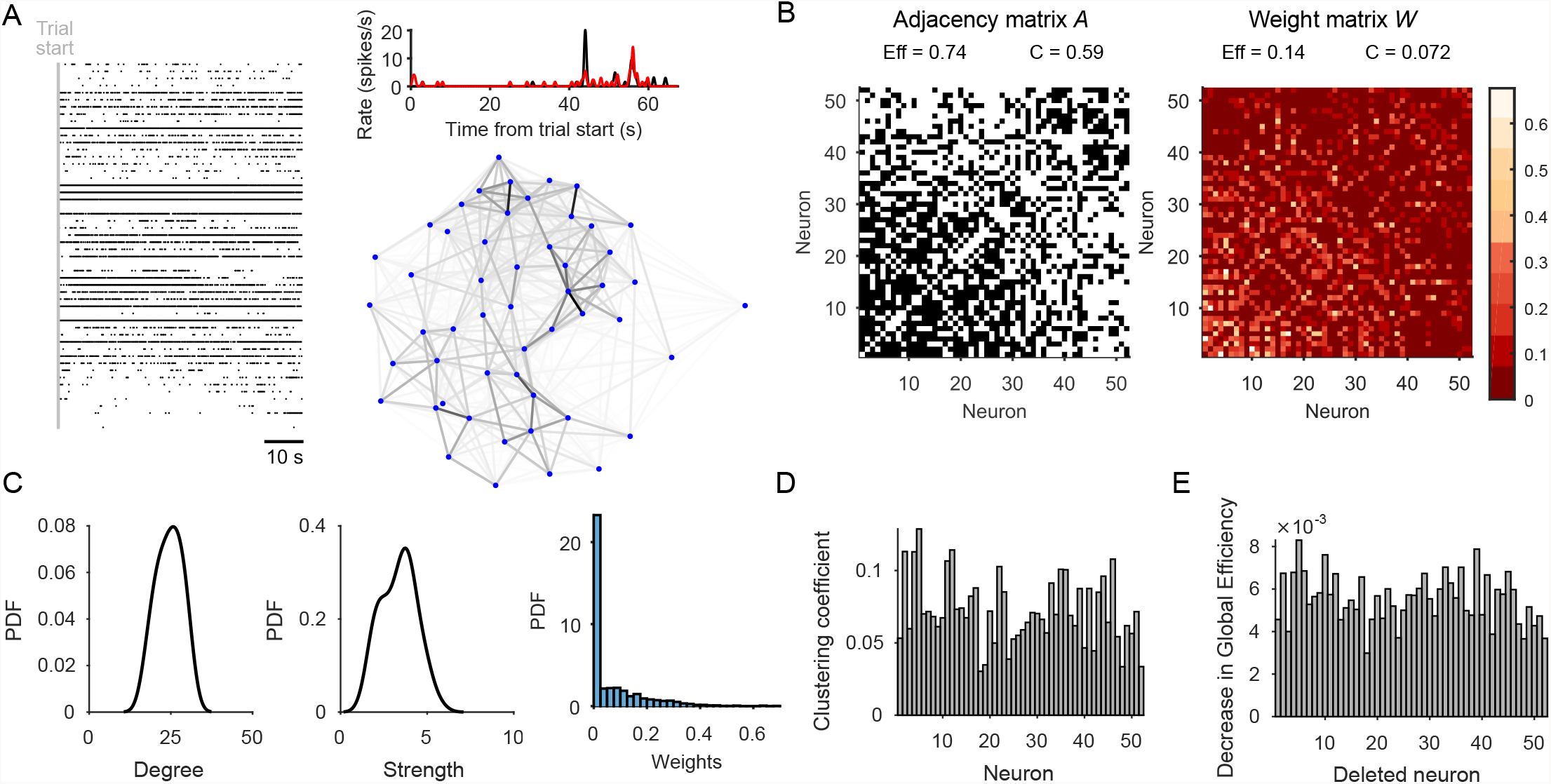
Quantifying neural population dynamics using network science. A) Schematic of turning neural activity time-series into a network. Left: a raster plot of 52 simultaneously recorded neurons in rat medial prefrontal cortex, during a single trial of a maze navigation task. Right: the corresponding network representation: nodes are neurons, links indicate pairwise interactions, and their grey-scale indicates the strength of interaction. Top: Interactions here are rectified Pearson’s *R* (setting *R* < 0 to 0) between pairs of spike-trains convolved with a Gaussian (*σ* = 250 ms); two example convolved trains are plotted here. B) Representations of the network in panel A: the adjacency matrix describes the presence (black) or absence (white) of links; the weight matrix describes the strengths of those links. Neurons are ranked by total link strength in descending order. Above each we give the global efficiency (*E f f)* and average clustering coefficient (*C*), respectively measuring the ease of getting from one node to another, and the density of links in the neighbourhood of one node. C) Distributions of node degree (total number of links per node), node strength (total weight of links per node), and link strength for the network in panel A. D) Network clustering fingerprint. A histogram of the weighted clustering coefficient for each neuron, measuring the ratio of weighted triangles to weighted triples in which that neuron participates: the higher the ratio, the more strongly connected is the neighbourhood of that neuron. Some neurons (e.g. 2, 5) have strongly connected neighbourhoods, implying a local group of correlated neurons. E) Network efficiency fingerprint, given by the decrease in the network’s global efficiency after deleting each neuron in turn. Neurons that strongly decrease the efficiency (e.g. 3) are potential network hubs, mediating interactions between many neurons.

We are free to choose any measure of pairwise interaction we like; and indeed that choice depends on what questions we want to ask of the data. Typical choices include cosine similarity or a rectified correlation coefficient, as these linear measures are familiar, easy to interpret, and not data-intensive. But with sufficient data we could also use non-linear measurements of interaction including forms of mutual information (Bettencourt, Stephens, Ham, & Gross, 2007; Singh & Lesica, 2010) and transfer entropy (Nigam et al., 2016; Schreiber, 2000; Thivierge, 2014). We could fit an Ising model, so estimating “direct” interactions while factoring out other inputs (S. Yu, Huang, Singer, & Nikolic, 2008). We could even fit a model to each neuron for the generation of its activity time-series, such as a generalised linear model (Pillow et al., 2008; Truccolo, Eden, Fellows, Donoghue, & Brown, 2005), and use the fitted weights of the inputs from all other neurons as the interaction values in a network (Gerhard, Pipa, Lima, Neuenschwander, & Gerstner, 2011). In addition, there is a large selection of interaction measures specific for spike-trains (e.g. Lyttle & Fellous, 2011; van Rossum, 2001; Victor & Purpura, 1996), whose use in defining interaction networks has yet to be well explored. And we should always be mindful that measures of pairwise interaction alone cannot distinguish between correlations caused by common input from unrecorded neurons and correlations caused by some direct contact between the recorded neurons.

Whatever measure of interaction we use, the important distinction is between whether the interaction measurement is undirected (e.g. the correlation coefficient) or directed (e.g. transfer entropy), and so whether we end up with an undirected or directed network as a result (throughout this paper I consider only symmetric measures of interaction, and hence undirected networks). And we end up with a weighted network (Newman, 2004). While much of network science, and its use in neuroscience, is focussed on binary networks whose links indicate only whether or not an interaction between two nodes exists, any measurement of interaction gives us a weight for each link (Figure 1B). Thresholding the weights to construct a binary network inevitably loses information (Humphries, 2011; Zanin et al., 2012). Consequently, multi-neuron recording data are best captured in a weighted network.

This weighted network of interactions between neurons need not map to any physical network of connections between neurons. The synaptic connections between neurons in a circuit shape and constrain the dynamics of those neurons, which we capture as population activity in multi-neuron recordings. But interactions can change independently of the physical network, both because the firing of a single neuron requires inputs from many other neurons, and because physical connections can be modulated on fast time-scales, such as short-term plasticity temporarily enhancing or depressing the strength of a synapse. Nonetheless, because physical connections between neurons constrain their dynamics, so sustained changes in interactions on time-scales of minutes and hours are evidence of some physical change to the underlying circuit (Baeg et al., 2007; Carrillo-Reid, Yang, Bando, Peterka, & Yuste, 2016; Grewe et al., 2017; Laubach, Wessberg, & Nicolelis, 2000; Yamada et al., 2017).

The use of network science to describe interactions between neural elements has been growing in cognitive neuroscience for a decade, and widely used to analyse EEG, MEG, and fMRI time-series data (Achard, Salvador, Whitcher, Suckling, & Bullmore, 2006; Bassett & Bullmore, 2016; Bullmore & Sporns, 2009). Neuroimaging has long used the unfortunate term “functional networks”, with its connotations of causality and purpose, to describe the network of pairwise correlations between time-series of neural activity. To avoid any semantic confusion, and distinguish the networks of interactions from the underlying physical network, I will describe the network of single neuron interactions here as a “dynamical” network.

What can we do with such dynamical networks of neurons? In the following I show how with them we can quantify circuit-wide changes following perturbations and manipulations; we can track changes in dynamics over time; and we can quantitatively define qualitative theories of computational concepts.

Efficiency:

Reciprocal of the mean shortest path length between all pairs of nodes; path lengths are weighted. The higher the efficiency, the shorter the average path between a pair of nodes.

Small-world network:

A network with both high clustering of nodes and high efficiency.

## CAPTURING POPULATION DYNAMICS AND THEIR CHANGES BY MANIPULATIONS

Applying network science to large-scale recordings of neural systems allows us to capture their complex dynamics in a compact form. The existing toolbox of network science gives us a plethora of options for quantifying the structure of a dynamical network. We may simply quantify its degree and strength distributions (Figure 1C), revealing dominant neurons (Dann, Michaels, Schaffelhofer, & Scherberger, 2016; Nigam et al., 2016). We can assess the local clustering of the dynamical network, the proportion of a neuron’s linked neighbours that are also strongly linked to each other (Watts & Strogatz, 1998; Figure 1D), revealing the locking of dynamics among neurons (Bettencourt et al., 2007; Sadovsky & MacLean, 2013). We can compute the efficiency of a network (Latora & Marchiori, 2001), a measure of how easily a network can be traversed (Figure 1E), revealing how cohesive the dynamics of a population are - the higher the efficiency, the more structured the interactions amongst the entire population (Thivierge, 2014). We may define structural measures relative to a null model, such as quantifying how much of a small-world the dynamical network is (Dann et al., 2016; Gerhard et al., 2011; S. Yu et al., 2008). Our choice of quantifying measures depends on the aspects of dynamics we are most interested in capturing.

Having compactly described the dynamics, we are well-placed to then characterise the effects of manipulating that system. Manipulations of a neural system will likely cause system-wide changes in its dynamics. Such changes may be the fast, acute effect of opto-genetic stimulation (Boyden, 2015; Deisseroth, 2015; Miesenböck, 2009); the sluggish but acute effects of drugs (Vincent, Tauskela, Mealing, & Thivierge, 2013); or the chronic effects of neurological damage (Otchy et al., 2015). All these manipulations potentially change the interactions between neurons, disrupting normal computation. By comparing the dynamical networks before and after the manipulation, one could easily capture the changes in the relationships between neurons.

There have been few studies examining this idea. Srinivas, Jain, Saurav, and Sikdar (2007) used dynamical networks to quantify the changes to network-wide activity in hippocampus caused by the glutamate-injury model of epilepsy, suggesting a dramatic drop in network clustering in the epilepsy model. Vincent et al. (2013) used dynamical networks to quantify the potential neuroprotective effects of drug pre-conditioning in rat cortex *in vitro*, finding increased clustering and increased efficiency in the network over days, implying the drugs enriched the synaptic connections between groups of neurons. Quantifying manipulations using network science is an under-explored application, rich in potential.

## TRACKING THE EVOLUTION OF DYNAMICS

Neural activity is inherently non-stationary, with population activity moving between different states on a range of time-scales, from shifting global dynamics on time-scales of seconds (Zagha & McCormick, 2014), to changes wrought by learning on time-scales of minutes and hours (Benchenane et al., 2010; Huber et al., 2012). For a tractable understanding of these complex changes, ideally we would like a way describe the entire population’s dynamics with as few parameters as possible. A recent example of such an approach is population coupling, the correlation over time between a single neuron’s firing rate and the population average rate (Okun et al., 2015). But with dynamical networks we can use the same set of tools above, and more, to easily track changes to the population activity in time.

Figure 2 illustrates the idea of tracking non-stationary activity with data from a study by Peyrache, Khamassi, Benchenane, Wiener, and Battaglia (2009). Rats were required to learn rules in a Y-maze to obtain reward. I use here a single session in which a rat learned the rule “go to the cued arm” (Figure 2A); 52 simultaneously recorded neurons from medial prefrontal cortex were active in every trial of this session. As the rat learned the rule in this session, and activity in medial prefrontal cortex is known to represent changes in behavioural strategy (Durstewitz, Vittoz, Floresco, & Seamans, 2010; Karlsson, Tervo, & Karpova, 2012; Powell & Redish, 2016), we might reasonably expect the population activity to evolve during rule-learning. Visualising trial-by-trial changes using dynamical networks (built as in Figure 1A) shows a stabilisation of the interactions between neurons over trials (Figure 2B). Quantifying this by correlating weight matrices on consecutive trials (Figure 2C), confirms there was a rapid stabilisation of neuron interactions at the start of this learning session. Plotting the total weight or total number of links in the network over trials (Figure 2D) shows that this stabilisation of the dynamical network was not a simple consequence of a global stabilisation of the interactions between neurons. These analyses thus track potentially learning-induced changes in the population activity of prefrontal cortex.

**Figure 2.**
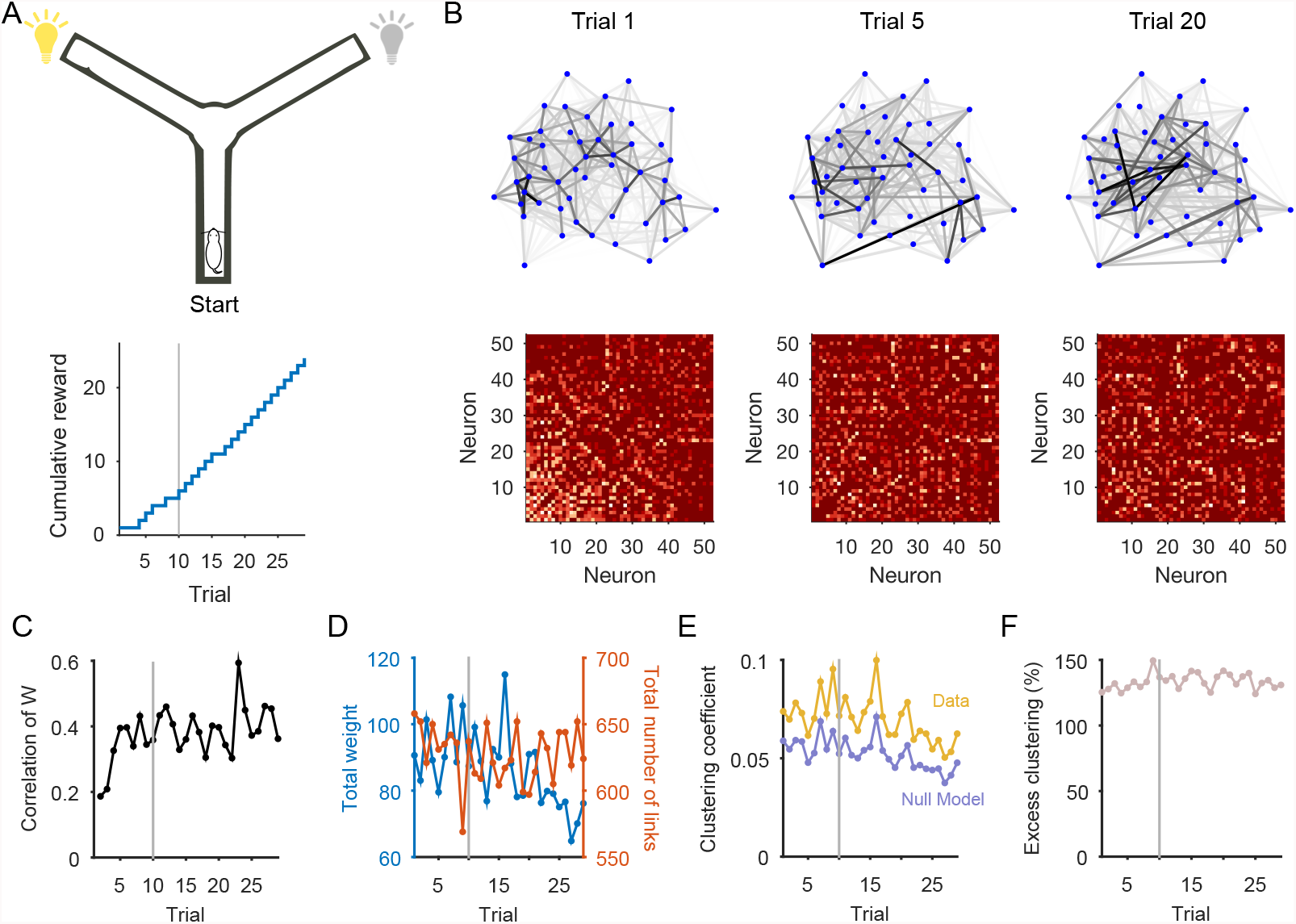
Tracking changes in neural population dynamics using network science. A) Recordings examined here are from one behavioural session of a Y-maze learning task. For this session, the rat had to reach the end of the randomly-cued arm to receive reward (schematic, top). This session showed evidence of behavioural learning (bottom), with a sustained increase in reward accumulation after trial10 (grey line). A trial lasted typically 70 s, running from the rat leaving the start position through reaching the arm end and returning to the start position to initiate the next trial. The population activity from a single trial is shown in Fig. 1A. B) Dynamical networks from trials 1, 5 and 20 of that session. The top row plots the networks, with nodes as neurons and greyscale links indicating the strength of pairwise interaction. The bottom row plots the corresponding weight matrix (ordered by total node strength in trial1 throughout). The networks show a clear re-organisation of interactions between neurons during learning. C) Tracking network stability. The correlation between the weight matrix W at trial *t* and at trial *t* − 1. The dynamical network rapidly increased in similarity over the first few trials. Grey line: behavioural learning trial. D) Changes in total weight (red) and total number of links (blue) over trials. E) Clustering coefficient of the weighted network (‘Data’) on each trial; compared to the mean clustering coefficient over 20 null model weighted networks per trial (‘Null Model’). F) Excess clustering in the data compared to the null model on each trial (data in panel E expressed as a ratio: 100 x Cd a ta *I* CmodeJ). The va riation across trials in the data is well-accounted for by the null model, suggesting the average local clustering did not change over learning.

We can also use these data to illustrate the benefits we accrue from the null models in network science. These models define the space of possible networks obtained by some stochastic process. Classically, the null model of choice was the Erdos-Renyi random network, which assumes a uniform probability of a link falling between any pair of nodes. As few if any real-world networks can be described this way, more detailed null models are now available. One common example is the configuration model (Chung & Lu, 2002; Fosdick, Larremore, Nishimura, & Ugander, 2016), in which we assume connections between nodes are made proportional to the number of links they already have. This model, applied to neural time-series, is a null model for testing whether the existence of interactions between a pair of neurons is simply a result of those neurons having many interactions. Other null model networks include the exponential random graph model (Robins, Pattisona, Kalisha, & Lushera, 2007), or the stochastic block model and its variants (Newman & Martin, 2014). In general, network null models allow us to test whether features of our dynamical networks exceed those expected by stochastic variation alone.

Clustering coefficient:

Ratio of weighted triangles to weighted triples - incomplete triangles - in the network.

Motifs:

A specific pattern of connections between a small number of nodes, which includes at least one connection for every node. For example, in an undirected network, for 4 nodes there are 5 possible motifs.

We use the example of determining whether or not there is a change in the clustering of interactions between neurons over this example learning session. Figure 2E plots the average clustering coefficient for the dynamical networks, and we can see that it varies across trials. We can compare this to a suitable null model; here I use a null model that conserves node strength, but randomly re-assigns the set of weights between nodes (Rubinov & Sporns, 2011). Plotting the average clustering coefficient for this null model on each trial shows that the clustering in the data-derived dynamical networks is well in excess of that predicted by the null model: the interactions between groups of three neurons are more dense than predicted by just their total interactions with all neurons.

But the null model also shows that the average local clustering does not change over learning. The ratio of the data and model clustering coefficients is approximately constant (Figure 2F), showing that trial-by-trial variation in clustering is largely accounted for by variations in the overall interactions between neurons (one source of these might be finite-size effects in estimating the interactions on trials of different durations). So we can conclude that changes over behavioural learning in this population of neurons reflected a local reorganisation (Figure 2B) and stabilisation (Figure 2C) of interactions, but which did not change the population-wide distribution of clustering.

The rich potential for tracking dynamics with the readily-available metrics of network science has not yet been tapped. As just demonstrated, with dynamical networks we can track trial-by-trial or event-by-event changes in population dynamics. For long recordings of spontaneous activity, building dynamical networks in time-windows slid over the recorded data allows us to track hidden shifts underlying global dynamics (Humphries, 2011). On slower time-scales, we can track changes during development of neural systems, either using *ex-vivo* slices (Dehorter et al., 2011) or *in vitro* cultures (Downes et al., 2012; M. S. Schroeter, Charlesworth, Kitzbichler, Paulsen, & Bullmore, 2015). These studies of development have all shown how maturing neuronal networks move from seemingly randomly-distributed interactions between neurons to a structured set of interactions, potentially driven by changes to the underlying connections between them.

Other tools from network science could be readily re-purposed to track neural population dynamics. The growing field of network comparison uses distributions of network properties to classify networks (Guimera, Sales-Pardo, & Amaral, 2007; Onnela et al., 2012; Przulj, 2007; Wegner, Ospina-Forero, Gaunt, Deane, & Reinert, 2017). A particularly promising basis for comparison is the distributions of motifs (or graphlets) in the networks (Przulj, 2007). Re-purposed to track changes in dynamical networks, by comparing motif distributions between time-points, these would provide tangible evidence of changes to the information flow in a neural system.

Ongoing developments in temporal networks (Holme, 2015) and network-based approaches to change-point detection algorithms (Barnett & Onnela, 2016; Darst et al., 2016; Peel & Clauset, 2014) also promise powerful yet tractable ways to track neural population dynamics. Temporal networks in particular offer a ranges of formalisms for tracking changes through time (Holme, 2015). In one approach, interaction networks for each slice of time are coupled by links between the same node in adjacent time-slices; this allows testing for how groups of nodes evolve over time, constrained by their groups in each slice of time (Bassett et al., 2011; Mucha, Richardson, Macon, Porter, & Onnela, 2010). A range of null models are available for testing the evolution of networks in this time-slice representation (Bassett et al., 2013). But such a representation requires coarse-graining of time to capture the interactions between all nodes in each time-slice. An alternative approach is to define a network per small time-step, comprising just the interactions that exist at each time-step (Holme, 2015; Thompson, Brantefors, & Fransson, 2016), and then introduce the idea of reachability: that one node is reachable from another if they both link to an intermediate node on different time-steps. With this representation, standard network measures such as path-lengths, clustering, and motifs can be easily generalised to include time (Thompson et al., 2016). Thus, a network description of multi-neuron activity need not just be a frozen snapshot of interactions, but can be extended to account for changes in time.

## NETWORK THEORY QUANTITATIVELY DEFINES COMPUTATIONAL CONCEPTS OF NEURAL POPULATIONS

The mathematical framework of networks can also provide precise quantitative definitions of important but qualitative theories about neural populations. A striking example is the theory of neural ensembles (Harris, 2005). An ensemble is qualitatively defined as a set of neurons that are consistently co-active (Harris, 2005), thereby indicating they code or compute the same thing. This qualitative definition leaves open key quantitative questions: what defines co-active, and what defines consistent?

The network science concept of modularity provides answers to these questions. Many networks are modular, organised into distinct groups: social networks of friendship groups, or collaboration networks of scientists. Consequently, the problem of finding modules within networks in an unsupervised way is an extraordinarily fecund research field (Fortunato & Hric, 2016). Most approaches to finding modules are based on the idea of finding the division of the network that maximises its modularity *Q* = {number of links within a module} - {expected number of such links} (Newman, 2006). Maximising *Q* thus finds a division of a network in which the modules are densely linked within themselves, and weakly linked between them.

Applied to dynamical networks, modularity defines neural ensembles (Billeh, Schaub, Anastassiou, Barahona, & Koch, 2014; Bruno et al., 2015; Humphries, 2011): groups of neurons that are more co-active with each other than with any other neurons in the population, given the choice of pairwise interaction used. Figure 3 demonstrates this idea using an example recording of 94 neurons from the motor circuit of the sea-slug *Aplysia* during fictive locomotion (Bruno et al., 2015). The weight matrix and network view in Figure 3A clearly indicate some structure within the dynamical network. Applying an unsupervised module-detection algorithm finds a high modularity division of the dynamical network (Figure 3B). When we plot the 94 spike-trains grouped by their modules in the dynamical network, the presence of multiple ensembles is clear (Figure 3C).

**Figure 3.**
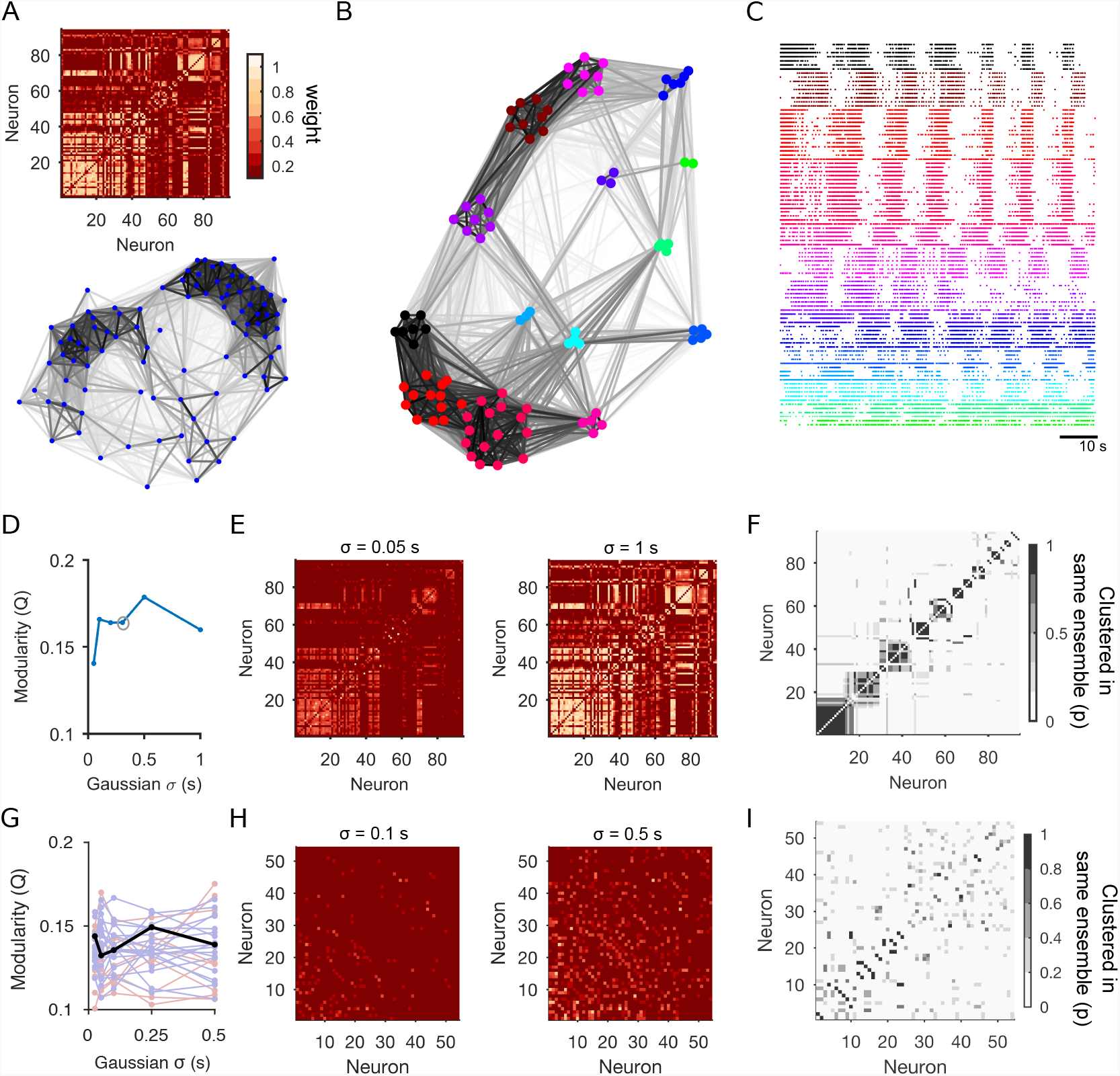
Defining and detecting neural ensembles using network science.

With this modularity-based approach, we can also easily check how robust these ensembles are to the choice of time-scale of co-activity. When computing pairwise interactions, we often have a choice of temporal precision, such as bin-size or Gaussian width (Figure 1A): choosing small values emphasises spike-time precision; large values emphasise co-varying firing rates. As shown in Figure 3D, we can also use *Q* to look for time-scales at which the population dynamics are most structured (Humphries, 2011): this view suggests a clear peak time-scale at which the ensembles are structured. Nonetheless, we can also see a consistent set of modules at all time-scales: the weight matrix *W* at the smallest and largest A dynamical network of population dynamics during crawling in *Aplysia.* The weight matrix (top) and network view (bottom) for a simultaneous recording of 94 neurons during 90 seconds from the initiation of crawling (from the experimental protocol of Bruno et al., 2015). Weights are rectified Pearson’s *R* between pairs of neurons convolved with a Gaussian of *σ* = 0.306 s (using the median inter-spike interval of the recording as an initial guide to time-scale, as in Bruno et al., 2015). B) Modules within the dynamical network. Coloured nodes indicate different modules found within the dynamical network using an unsupervised consensus module-detection algorithm (Bruno et al., 2015). Placement of the modules reflects the similarity between them (Traud et al., 2009). C) Raster plot of the corresponding spike-trains, grouped according to the modules in panel B. The detection of multiple neural ensembles is evident. D) Dependence of the modular structure on the time-scale of correlation. Smaller Gaussian *CT* detects precise spike-timing; larger *fT* detects co-variation in firing rates. Circle: time-scale used in panels A-C. E) Weight matrices for the smallest and largest time-scale used for the Gaussian convolution. Neurons are plotted in descending order of total weight in the shorter time-scale. F) Stability of modules over time-scales. The confusion matrix showing for each pair of neurons the proportion of time-scales for which that pair was placed in the same module. The majority of neuron pairs were placed in the same module at every time-scale. G)-I) Comparable analysis for the medial prefrontal cortex data. G) Dependence of Q on the time-scale of correlation, for every trial in one session (from Fig. 2). Black: learning trial; red: pre-learning trial; blue: post-learning trial. H) As for panel E, for the learning trial of the medial prefrontal cortex data. I) As for panel F, for the learning trial.

Gaussian width are similar (Figure 3E); and the majority of neurons are placed in the same group at every time-scale (Figure 3F). Modularity not only defines ensembles, but also lets us quantify their time-scales and find consistent structure across time-scales.

The complexity of the population activity will determine whether a consistent set of ensembles appears across time-scales, or whether there are different ensembles at different time-scales (see Humphries, 2011, for more examples). We can see this when running the same module-detection analysis on a session from the medial prefrontal cortex data (Figure 3G-I). For this cortical data there are modules present at every time-scale, but no consistent time-scale at which the neural activity is most structured (Figure 3G-H). Consequently, there is not a consistent set of modules across time-scales (Figure 3I).

Such multi-scale structure is potentially a consequence of the order-of-magnitude distribution in firing rates (Dann et al., 2016; Wohrer et al., 2013), for which more work is needed on suitable measures of interaction. It may also indicate that some neurons are members of more than one ensemble, which are active at different times during the recording. Consequently, these neurons’ correlations with others will depend on the time-scale examined. Examining the detected modules for nodes that participate in more than one module (Guimera & Amaral, 2005; Guimera et al., 2007) may reveal these shared neurons. Clearly, such multi-scale structure means that tracking changes in the structure of population activity should be done at a range of time-scales, and comparisons made based on similar time-scales.

As a final step, we can now quantitatively define a Hebbian cell assembly (Holtmaat & Caroni, 2016). By definition, a cell assembly is an ensemble of neurons that become coactive because of changes to synaptic connections into and between them during learning (Carrillo-Reid et al., 2016). Thus, by combining the ideas of tracking dynamical networks and of module detection, we can test for the formation of assemblies: if we find dynamical network modules that appear during the course of learning, then we have identified potential cell assemblies.

### OUTLOOK

The dynamics of neural populations are emergent properties of the wiring within their microcircuits. We can of course use network science to describe physical networks of the microcircuit too (Humphries, Gurney, & Prescott, 2006; Lee et al., 2016; M. Schroeter, Paulsen, & Bullmore, 2017), gaining insight into the mapping from wiring to dynamics. But dynamical networks need not map to any circuit. Indeed while dynamical networks are constrained by their underlying physical connections, they can change faster than their corresponding physical networks. A clear example is with the actions of neuromodulators - these can increase or decrease the effective strength of connections between neurons and the responsiveness of individual neurons (Nadim & Bucher, 2014), so changing the dynamical network without changing the underlying physical network. More broadly, rapid, global changes in brain state can shift the dynamics of a neural population (Zagha & McCormick, 2014). Thus, dynamical networks describing the simultaneous activity of multiple neurons capture the moment-to-moment changes in population dynamics.

There are of course other analysis frameworks for visualising and describing the activity of large neural populations. The detection of neural ensembles is an unsupervised clustering problem, for which a number of neuroscience-specific solutions exist (Feldt, Waddell, Hetrick, Berke, & Zochowski, 2009; Fellous, Tiesinga, Thomas, & Sejnowski, 2004; Lopes-dos Santos, Conde-Ocazionez, Nicolelis, Ribeiro, & Tort, 2011; Russo & Durstewitz, 2017). Some advantages of network science here are that the detection of ensembles is but one application of the same representation of the population activity; that a range of null models is available for testing hypotheses of clustering; and that the limitations of module-detection are well established, allowing comparatively safe interpretation of the results (Fortunato & Hric, 2016; Good, de Montjoye, & Clauset, 2010). More generally, analyses of neural population recordings have used dimension reduction approaches in order to visualise and describe the dynamics of the population (Cunningham & Yu, 2014; Pang, Lansdell, & Fairhall, 2016). As discussed in Box 1, both network and dimension-reduction approaches offer powerful, complementary views of complex neural dynamics.

#### Box 1. Networks and dimension-reduction approaches

Dimension reduction approaches to neural population recordings aim to find a compact description of the population’s activity using many fewer variables than neurons (Pang et al., 2016). Typical approaches include principal components analysis (PCA) and factor analysis, both of which aim to find a small set of dimensions in which the population activity can be described with minimal loss of information (Ahrens et al., 2012; Bartho, Curto, Luczak, Marguet, & Harris, 2009; Briggman et al., 2005; Bruno et al., 2015; Kato et al., 2015; Levi, Varona, Arshavsky, Rabinovich, & Selverston, 2005; Mazor & Laurent, 2005; Wohrer et al., 2013). More complex variants of these standard approaches can cope with widely-varying time-scales in cortical activity (B. M. Yu et al., 2009), or aim to decompose multiplexed encodings of stimulus variables by the population’s activity into different dimensions (Kobak et al., 2016).

Both network and standard dimension reduction approaches have in common the starting point of a pairwise interaction matrix. PCA, for example, traditionally uses the covariance matrix as its starting point. Consequently, both approaches assume that the relationships between neurons are static over the duration of the data from which the matrix is constructed. (This assumption is also true for dimension reduction methods that fit generative models, such as independent component analysis or Gaussian Process Factor Analysis (B. M. Yu et al., 2009), as fitting the model also assumes stationarity in the model’s parameters over the duration of the data).

Where the approaches diverge is in their advantages and limitations. Dimension reduction approaches offer the advantage of easy visualisation of the trajectories of the population activity over time. This in turn allows for potentially strong qualitative conclusions, either about the conditions under which the trajectories differ – such as in encoding different stimuli (Kobak et al., 2016; Mazor & Laurent, 2005) or making different decisions (Briggman et al., 2005; Harvey, Coen, & Tank, 2012) – or about the different states repeatedly visited by the population during movement (Ahrens et al., 2012; Kato et al., 2015; B. M. Yu et al., 2009). By contrast, there are not yet well-established ways of drawing quantitative conclusions from standard dimension reduction approaches, nor of how to track changes in the population dynamics over time, such as through learning. Further, while reducing the dimensions down to just those accounting for a high proportion of the variance (or similar) in the population activity can remove noise, it also risks removing some of the higher-dimensional, and potentially informative, dynamics in the population. Finally, to date, most applications of dimension reduction approaches have been based on just the pairwise covariance or correlation coefficient.

As I have demonstrated here, network-based approaches take a different slant on simplifying complex dynamics. The network description maintains a representation of every neuron, and so potentially captures all dynamical relationships that might be removed by dimension reduction. It is simple to use any measure of pairwise interaction, without changing the analysis. Quantitative analyses of either static (Figure 1) or changing (Figure 2) population activity is captured in simple, compact variables. And we have access to a range of null models for testing the existence of meaningful interactions between neurons and changes to those interactions. However, interpreting some of these quantifying variables, such as efficiency, in terms of neural activity is not straightforward. And it is not obvious how to visualise trial-by-trial population activity, nor how to draw qualitative conclusions about different trajectories or states of the activity. Consequently, combining both network and dimension-reduction approaches could offer complementary insights into a neural population’s dynamics (Bruno et al., 2015).

One motivation for turning to network science as a toolbox for systems neuroscience is rooted in the extraordinarily rapid advances in recording technology, now scaling to hundreds or thousands of simultaneously recorded neurons (Stevenson & Kording, 2011). Capturing whole nervous systems of even moderately complex animal models will require scaling by further orders of magnitude (Ahrens et al., 2012; Lemon et al., 2015). And here is where network science has its most striking advantage: these tools have been developed to address social and technological networks of millions of nodes or more, so easily scale to systems neuroscience problems now and in the foreseeable future.

This is not a one-way street. Systems neuroscience poses new challenges for network science. Most studies in network science concern a handful of static or slowly changing data networks. Neural populations have non-stationary dynamics, that change rapidly compared to the temporal resolution of our recordings. And systems neuroscience analysis requires quantitatively comparing multiple defined networks within and between brain regions, within and between animals, and across experimental conditions - stimuli, decisions, and other external changes. More work is needed, for example, on appropriate null models for weighted networks (Palowitch, Bhamidi, & Nobel, 2016; Rubinov & Sporns, 2011); and on appropriate ways to regularise such networks, in order to separate true interactions from stochastic noise (MacMahon & Garlaschelli, 2015). Bringing network science to bear on challenges in systems neuroscience will thus create a fertile meeting of minds.

## SUPPORTIVE INFORMATION

Visualisations and analyses here drew on a range of open-source MATLAB (Mathworks, NA) toolboxes:

- Brain Connectivity Toolbox (Rubinov & Sporns, 2010): https://sites.google.com/site/bctnet/
- Network visualisations used the MATLAB code of Traud et al. (2009), available here: http://netwiki.amath.unc.edu/VisComms. This also the needs MatlabBGL library: http://uk.mathworks.com/matlabcentral/fileexchange/10922-matlabbgl. Mac OSX 64-bit users will need this version: https://dgleich.wordpress.com/2010/07/08/matlabbgl-osx-64-bit/
- Spike-Train Communities Toolbox (Bruno et al., 2015; Humphries, 2011): implementing unsupervised consensus algorithms for module-detection https://github.com/mdhumphries/SpikeTrainCommunitiesToolBox

## ACKNOWLEDGMENTS

I thank Silvia Maggi for reading a draft, Adrien Peyrache for permission to use the rat medial prefrontal cortex data, and Angela Bruno and Bill Frost for permission to use the *Aplysia* pedal ganglion data. This work was funded by a Medical Research Council Senior non-Clinical Fellowship (MR/J008648/1), and a Medical Research Council research grant (MR/P005659/1).

## REFERENCES

Achard, S., Salvador, R., Whitcher, B., Suckling, J., & Bullmore, E. (2006). A resilient, low-frequency, small-world human brain functional network with highly connected association cortical hubs. J Neurosci, 26(1), 63–72.

Ahrens, M. B., Li, J. M., Orger, M. B., Robson, D. N., Schier, A. F., Engert, F., & Portugues, R. (2012). Brain-wide neuronal dynamics during motor adaptation in zebrafish. Nature, 485, 471–477.

Baeg, E. H., Kim, Y. B., Kim, J., Ghim, J.-W., Kim, J. J., & Jung, M. W. (2007). Learning-induced enduring changes in functional connectivity among prefrontal cortical neurons. J Neurosci, 27, 909–918.

Barnett, I., & Onnela, J.-P. (2016). Change point detection in correlation networks. Scientific Reports, 6, 18893.

Bartho, P., Curto, C., Luczak, A., Marguet, S. L., & Harris, K. D. (2009). Population coding of tone stimuli in auditory cortex: dynamic rate vector analysis. Eur J Neurosci, 30(9), 1767–1778.

Bassett, D. S., & Bullmore, E. T. (2016). Small-world brain networks revisited. The Neuroscientist.

Bassett, D. S., Porter, M. A., Wymbs, N. F., Grafton, S. T., Carlson, J. M., & Mucha, P. J. (2013). Robust detection of dynamic community structure in networks. Chaos, 23, 013142.

Bassett, D. S., Wymbs, N. F., Porter, M. A., Mucha, P. J., Carlson, J. M., & Grafton, S. T. (2011). Dynamic reconfiguration of human brain networks during learning. Proceedings of the National Academy of Sciences of the United States of America, 108, 7641–7646.

Benchenane, K., Peyrache, A., Khamassi, M., Tierney, P. L., Gioanni, Y., Battaglia, F. P., & Wiener, S. I. (2010). Coherent theta oscillations and reorganization of spike timing in the hippocampal-prefrontal network upon learning. Neuron, 66(6), 921–936.

Bettencourt, L. M. A., Stephens, G. J., Ham, M. I., & Gross, G. W. (2007). Functional structure of cortical neuronal networks grown in vitro. Phys Rev E, 75(2 Pt 1), 021915.

Billeh, Y. N., Schaub, M. T., Anastassiou, C. A., Barahona, M., & Koch, C. (2014). Revealing cell assemblies at multiple levels of granularity. J Neurosci Methods, 236, 92–106.

Boyden, E. S. (2015). Optogenetics and the future of neuroscience. Nat Neurosci, 18, 1200–1201.

Briggman, K. L., Abarbanel, H. D. I., & Kristan, W.Jr. (2005). Optical imaging of neuronal populations during decision-making. Science, 307(5711).

Bruno, A. M., Frost, W. N., & Humphries, M. D. (2015). Modular deconstruction reveals the dynamical and physical building blocks of a locomotion motor program. Neuron, 86, 304–318.

Bullmore, E., & Sporns, O. (2009). Complex brain networks: graph theoretical analysis of structural and functional systems. Nat Rev Neurosci, 10, 186–198.

Buzsáki, G. (2004). Large-scale recording of neuronal ensembles. Nat Neurosci, 7, 446–451.

Carrillo-Reid, L., Yang, W., Bando, Y., Peterka, D. S., & Yuste, R. (2016). Imprinting and recalling cortical ensembles. Science, 353, 691–694.

Chen, T.-W., Wardill, T. J., Sun, Y., Pulver, S. R., Renninger, S. L., Baohan, A.,… Kim, D. S. (2013). Ultrasensitive fluorescent proteins for imaging neuronal activity. Nature, 499, 295–300.

Chung, F., & Lu, L. (2002). Connected components in random graphs with given expected degree sequences. Annals of Combinatorics, 6, 124–145.

Cunningham, J. P., & Yu, B. M. (2014). Dimensionality reduction for large-scale neural recordings. Nat Neurosci, 17, 1500–1509.

Dann, B., Michaels, J. A., Schaffelhofer, S., & Scherberger, H. (2016). Uniting functional network topology and oscillations in the fronto-parietal single unit network of behaving primates. eLife, 5, e15719.

Darst, R. K., Granell, C., Arenas, A., Gó mez, S., Saramki, J., & Fortunato, S. (2016). Detection of timescales in evolving complex systems. Scientific Reports, 6, 39713.

Dehorter, N., Michel, F., Marissal, T., Rotrou, Y., Matrot, B., Lopez, C.,… Hammond, C. (2011). Onset of pup locomotion coincides with loss of NR2C/D-mediated corticostriatal EPSCs and dampening of striatal network immature activity. Front Cell Neurosci, 5, 24.

Deisseroth, K. (2015). Optogenetics: 10 years of microbial opsins in neuroscience. Nat Neurosci, 18(9), 1213–1225.

Downes, J. H., Hammond, M. W., Xydas, D., Spencer, M. C., Becerra, V. M., Warwick, K.,… Nasuto, S. J. (2012). Emergence of a small-world functional network in cultured neurons. PLoS Comput Biol, 8(5), e1002522.

Dupre, C., & Yuste, R. (2017). Non-overlapping neural networks in Hydra vulgaris. Current Biology, In press.

Durstewitz, D., Vittoz, N. M., Floresco, S. B., & Seamans, J. K. (2010). Abrupt transitions between prefrontal neural ensemble states accompany behavioral transitions during rule learning. Neuron, 66, 438–448.

Feldt, S., Waddell, J., Hetrick, V. L., Berke, J. D., & Zochowski, M. (2009). Functional clustering algorithm for the analysis of dynamic network data. Phys Rev E, 79, 056104.

Fellous, J. M., Tiesinga, P. H., Thomas, P. J., & Sejnowski, T. J. (2004). Discovering spike patterns in neuronal responses. J Neurosci, 24(12), 2989–3001.

Fortunato, S., & Hric, D. (2016). Community detection in networks: A user guide. Physics Reports, 659, 1–44.

Fosdick, B. K., Larremore, D. B., Nishimura, J., & Ugander, J. (2016). Configuring random graph models with fixed degree sequences. arXiv, 1608.00607.

Frady, E. P., Kapoor, A., Horvitz, E., & Kristan, W. B.Jr. (2016). Scalable semisupervised functional neurocartography reveals canonical neurons in behavioral networks. Neural Comput, 28(8), 1453–1497.

Gerhard, F., Pipa, G., Lima, B., Neuenschwander, S., & Gerstner, W. (2011). Extraction of network topology from multi-electrode recordings: is there a small-world effect? Front Comput Neurosci, 5, 4.

Good, B. H., de Montjoye, Y.-A., & Clauset, A. (2010). Performance of modularity maximization in practical contexts. Phys Rev E, 81, 046106.

Grewe, B. F., Grndemann, J., Kitch, L. J., Lecoq, J. A., Parker, J. G., Marshall, J. D.,… Schnitzer, M. J. (2017). Neural ensemble dynamics underlying a long-term associative memory. Nature, 543, 670–675.

Guimera, R., & Amaral, L. A. N. (2005). Cartography of complex networks: modules and universal roles. J Stat Mech, P02001.

Guimera, R., Sales-Pardo, M., & Amaral, L. A. N. (2007). Classes of complex networks defined by role-to-role connectivity profiles. Nat Phys, 3(1), 63–69.

Harris, K. D. (2005). Neural signatures of cell assembly organization. Nat Rev Neurosci, 6, 399–407.

Harris, K. D., Quiroga, R. Q., Freeman, J., & Smith, S. L. (2016). Improving data quality in neuronal population recordings. Nat Neurosci, 19, 1165–1174.

Harvey, C. D., Coen, P., & Tank, D. W. (2012). Choice-specific sequences in parietal cortex during a virtual-navigation decision task. Nature, 484(7392), 62–68.

Holme, P. (2015). Modern temporal network theory: A colloquium. Eur Phys J B, 88, 234.

Holtmaat, A., & Caroni, P. (2016). Functional and structural underpinnings of neuronal assembly formation in learning. Nat Neurosci, 19, 1553–1562.

Huber, D., Gutnisky, D. A., Peron, S., O’Connor, D. H., Wiegert, J. S., Tian, L.,… Svoboda, K. (2012). Multiple dynamic representations in the motor cortex during sensorimotor learning. Nature, 484(7395), 473–478.

Humphries, M. D. (2011). Spike-train communities: finding groups of similar spike trains. J Neurosci, 31, 2321–2336.

Humphries, M. D., Gurney, K., & Prescott, T. J. (2006). The brainstem reticular formation is a small-world, not scale-free, network. Proc Roy Soc B. Biol Sci, 273, 503–511.

Jun, J. J., Mitelut, C., Lai, C., Gratiy, S., Anastassiou, C., & Harris, T. D. (2017). Real-time spike sorting platform for high-density extracellular probes with ground-truth validation and drift correction. bioRxiv, 101030.

Karlsson, M. P., Tervo, D. G. R., & Karpova, A. Y. (2012). Network resets in medial prefrontal cortex mark the onset of behavioral uncertainty. Science, 338, 135–139.

Kato, S., Kaplan, H. S., Schrödel, T., Skora, S., Lindsay, T. H., Yemini, E.,… Zimmer, M. (2015). Global brain dynamics embed the motor command sequence of caenorhabditis elegans. Cell, 163, 656–669.

Kobak, D., Brendel, W., Constantinidis, C., Feierstein, C. E., Kepecs, A., Mainen, Z. F.,… Machens, C. K. (2016). Demixed principal component analysis of neural population data. Elife, 5, e10989.

Latora, V., & Marchiori, M. (2001). Efficient behaviour of small-world networks. Phys Rev Lett, 87, 198701.

Laubach, M., Wessberg, J., & Nicolelis, M. A. (2000). Cortical ensemble activity increasingly predicts behaviour outcomes during learning of a motor task. Nature, 405(6786).

Lee, W.-C. A., Bonin, V., Reed, M., Graham, B. J., Hood, G., Glattfelder, K., & Reid, R. C. (2016). Anatomy and function of an excitatory network in the visual cortex. Nature, 532, 370–374.

Lemon, W. C., Pulver, S. R., Hö ckendorf, B., McDole, K., Branson, K., Freeman, J., & Keller, P. J. (2015). Whole-central nervous system functional imaging in larval Drosophila. Nat Commun, 6, 7924.

Levi, R., Varona, P., Arshavsky, Y. I., Rabinovich, M. I., & Selverston, A. I. (2005). The role of sensory network dynamics in generating a motor program. J Neurosci, 25(42), 9807–9815.

Lopes-dos Santos, V., Conde-Ocazionez, S., Nicolelis, M. A. L., Ribeiro, S. T., & Tort, A. B. L. (2011). Neuronal assembly detection and cell membership specification by principal component analysis. PLoS One, 6(6), e20996.

Lyttle, D., & Fellous, J.-M. (2011). A new similarity measure for spike trains: Sensitivity to bursts and periods of inhibition. J Neurosci Methods, 199(2), 296–309.

MacMahon, M., & Garlaschelli, D. (2015). Community detection for correlation matrices. Phys. Rev. X, 5, 021006.

Mazor, O., & Laurent, G. (2005). Transient dynamics versus fixed points in odor representations by locust antennal lobe projection neurons. Neuron, 48(4), 661–673.

Miesenböck, G. (2009). The optogenetic catechism. Science, 326(5951), 395–399.

Mucha, P. J., Richardson, T., Macon, K., Porter, M. A., & Onnela, J.-P. (2010). Community structure in time-dependent, multiscale, and multiplex networks. Science, 328(5980), 876–878.

Nadim, F., & Bucher, D. (2014). Neuromodulation of neurons and synapses. Curr Opin Neurobiol, 29, 48–56.

Newman, M. E. J. (2003). The structure and function of complex networks. SIAM Review, 45, 167–256.

Newman, M. E. J. (2004). Analysis of weighted networks. Phys Rev E, 70, 056131.

Newman, M. E. J. (2006). Finding community structure in networks using the eigenvectors of matrices. Phys Rev E, 74, 036104.

Newman, M. E. J., & Martin, T. (2014). Equitable random graphs. Phys Rev E, 90, 052824.

Nigam, S., Shimono, M., Ito, S., Yeh, F.-C., Timme, N., Myroshnychenko, M.,… Beggs, J. M. (2016). Rich-club organization in effective connectivity among cortical neurons. J Neurosci, 36, 670–684.

Okun, M., Steinmetz, N. A., Cossell, L., Iacaruso, M. F., Ko, H., Barth, P.,… Harris, K. D. (2015). Diverse coupling of neurons to populations in sensory cortex. Nature, 521, 511–515.

Onnela, J.-P., Fenn, D. J., Reid, S., Porter, M. A., Mucha, P. J., Fricker, M. D., & Jones, N. S. (2012). Taxonomies of networks from community structure. Phys Rev E, 86, 036104.

Otchy, T. M., Wolff, S. B. E., Rhee, J. Y., Pehlevan, C., Kawai, R., Kempf, A.,… Ölveczky, B. P. (2015). Acute off-target effects of neural circuit manipulations. Nature, 528, 358–363.

Palowitch, J., Bhamidi, S., & Nobel, A. B. (2016). The continuous configuration model: A null for community detection on weighted networks. arXiv, 1601.05630.

Pang, R., Lansdell, B. J., & Fairhall, A. L. (2016). Dimensionality reduction in neuroscience. Curr Biol, 26(14), R656–R660.

Peel, L., & Clauset, A. (2014). Detecting change points in the large-scale structure of evolving networks. arXiv, 1403.0989.

Peron, S., Chen, T.-W., & Svoboda, K. (2015). Comprehensive imaging of cortical networks. Curr Opin Neurobiol, 32, 115–123.

Peron, S. P., Freeman, J., Iyer, V., Guo, C., & Svoboda, K. (2015). A cellular resolution map of barrel cortex activity during tactile behavior. Neuron, 86, 783–799.

Peyrache, A., Khamassi, M., Benchenane, K., Wiener, S. I., & Battaglia, F. P. (2009). Replay of rule-learning related neural patterns in the prefrontal cortex during sleep. Nat Neurosci, 12, 916–926.

Pillow, J. W., Shlens, J., Paninski, L., Sher, A., Litke, A. M., Chichilnisky, E. J., & Simoncelli, E. P. (2008). Spatio-temporal correlations and visual signalling in a complete neuronal population. Nature, 454, 995–999.

Pouget, A., Beck, J. M., Ma, W. J., & Latham, P. E. (2013). Probabilistic brains: knowns and unknowns. Nat Neurosci, 16(9), 1170–1178.

Powell N. J., & Redish, A. D. (2016). Representational changes of latent strategies in rat medial prefrontal cortex precede changes in behaviour. Nat Commun, 7, 12830.

Przulj, N. (2007). Biological network comparison using graphlet degree distribution. Bioinformatics, 23, e177–e183.

Rieke, F., Warland, D., de Ruyter van Stevninck, R., & Bialek, W. (1999). Spikes: Exploring the neural code. Cambridge MA: MIT Press.

Robins, G., Pattisona, P., Kalisha, Y., & Lushera, D. (2007). An introduction to exponential random graph (p*) models for social networks. Social Networks, 29, 173–191.

Rubinov, M., & Sporns, O. (2010). Complex network measures of brain connectivity: uses and interpretations. Neuroimage, 52, 1059–1069.

Rubinov, M., & Sporns, O. (2011). Weight-conserving characterization of complex functional brain networks. Neuroimage, 56, 2068–2079.

Russo, E., & Durstewitz, D. (2017). Cell assemblies at multiple time scales with arbitrary lag constellations. eLife, 6, e19428.

Sadovsky, A. J., & MacLean, J. N. (2013). Scaling of topologically similar functional modules defines mouse primary auditory and somatosensory microcircuitry. J Neurosci, 33, 14048–60.

Schreiber, T. (2000). Measuring information transfer. Phys Rev Lett, 85, 461–464.

Schroeter, M., Paulsen, O., & Bullmore, E. T. (2017). Micro-connectomics: probing the organization of neuronal networks at the cellular scale. Nat Rev Neurosci, in press.

Schroeter, M. S., Charlesworth, P., Kitzbichler, M. G., Paulsen, O., & Bullmore, E. T. (2015). Emergence of rich-club topology and coordinated dynamics in development of hippocampal functional networks in vitro. J Neurosci, 35, 5459–5470.

Singh, A., & Lesica, N. A. (2010). Incremental mutual information: a new method for characterizing the strength and dynamics of connections in neuronal circuits. PLoS Comput Biol, 6, e1001035.

Srinivas, K. V., Jain, R., Saurav, S., & Sikdar, S. K. (2007). Small-world network topology of hippocampal neuronal network is lost, in an in vitro glutamate injury model of epilepsy. Eur J Neurosci, 25, 3276–3286.

Stevenson, I. H., & Kording, K. P. (2011). How advances in neural recording affect data analysis. Nat Neurosci, 14, 139–142.

Thivierge, J.-P. (2014). Scale-free and economical features of functional connectivity in neuronal networks. Phys Rev E, 90, 022721.

Thompson, W. H., Brantefors, P., & Fransson, P. (2016). From static to temporal network theory - applications to functional brain connectivity. bioRxiv, 096461.

Traud, A. L., Frost, C., Mucha, P. J.,, & Porter, M. A. (2009). Visualization of communities in networks. Chaos, 19, 041104.

Truccolo, W., Eden, U. T., Fellows, M. R., Donoghue, J. P., & Brown, E. N. (2005). A point process framework for relating neural spiking activity to spiking history, neural ensemble, and extrinsic covariate effects. J Neurophysiol, 93, 1074–1089.

van Rossum, M. C. (2001). A novel spike distance. Neural Comput, 13(4), 751–763.

Victor, J. D., & Purpura, K. P. (1996). Nature and precision of temporal coding in visual cortex: a metric-space analysis. J Neurophysiol, 76(2), 1310–1326.

Vincent, K., Tauskela, J. S., Mealing, G. A., & Thivierge, J.-P. (2013). Altered network communication following a neuroprotective drug treatment. PloS One, 8, e54478.

Watts, D. J., & Strogatz, S. H. (1998). Collective dynamics of ‘small-world’ networks. Nature, 393, 440–442.

Wegner, A. E., Ospina-Forero, L., Gaunt, R. E., Deane, C. M., & Reinert, G. (2017). Identifying networks with common organizational principles. arXiv, 1704.00387.

Wohrer, A., Humphries, M. D., & Machens, C. (2013). Population-wide distributions of neural activity during perceptual decision-making. Prog Neurobiol, 103, 156–193.

Yamada, Y., Bhaukaurally, K., Madarsz, T. J., Pouget, A., Rodriguez, I., & Carleton, A. (2017). Context- and output layer-dependent long-term ensemble plasticity in a sensory circuit. Neuron, 93, 1198–1212.e5.

Yu, B. M., Cunningham, J. P., Santhanam, G., Ryu, S. I., Shenoy, K. V., & Sahani, M. (2009). Gaussian-process factor analysis for low-dimensional single-trial analysis of neural population activity. J Neurophysiol, 102, 614–635.

Yu, S., Huang, D., Singer, W., & Nikolic, D. (2008). A small world of neuronal synchrony. Cereb Cortex, 18, 2891–2901.

Yuste, R. (2015). From the neuron doctrine to neural networks. Nat Rev Neurosci, 16, 487–497.

Zagha, E., & McCormick, D. A. (2014). Neural control of brain state. Curr Opin Neurobiol, 29, 178–186.

Zanin, M., Sousa, P., Papo, D., Bajo, R., Garca-Prieto, J., del Pozo, F.,… Boccaletti, S. (2012). Optimizing functional network representation of multivariate time series. Sci Rep, 2, 630.

